# High-dimensional multi-omics measured in controlled conditions are useful for maize platform and field trait predictions

**DOI:** 10.1101/2024.05.30.596567

**Authors:** Ali Baber, Huguenin-Bizot Bertrand, Laurent Maxime, Chaumont François, C Maistriaux Laurie, Nicolas Stéphane, Duborjal Hervé, Welcker Claude, Tardieu François, Mary-Huard Tristan, Moreau Laurence, Charcosset Alain, Runcie Daniel, Rincent Renaud

## Abstract

The effects of climate change in the form of drought, heat stress, and irregular seasonal changes threaten global crop production. The ability of multi-omics data, such as transcripts and proteins, to reflect a plant’s response to such climatic factors can be capitalized in prediction models to maximize crop improvement. Implementing multi-omics characterization in routine field evaluations is challenging due to high costs. It is, however, possible to do it on reference genotypes in controlled conditions. Using omics measured on a platform, we tested different multi-omics-based prediction approaches, with and without pedo-climatic data, using a high dimensional linear mixed model (MegaLMM) to predict genotypes for platform traits and agronomic field traits in a hybrid panel of 244 maize Dent lines crossed to a Flint tester. We considered two prediction scenarios: in the first one, new hybrids are predicted (CV1), and in the second one, partially observed hybrids are predicted (CV2). For both scenarios, all hybrids were characterized for omics on the platform. We observed that omics can predict both additive and non-additive genetic effects for the platform traits, resulting in much higher predictive abilities than GBLUP. This highlights their efficiency in capturing regulation processes in relation to the growth conditions. For the field traits, we observed that only the additive components of omics were useful and only slightly improved predictive abilities for predicting new hybrids (CV1, model MegaGAO) and for predicting partially observed hybrids (CV2, model GAOxW-BLUP) in comparison to GBLUP. We conclude that measuring the omics in the fields would be of considerable interest for predicting productivity, if the omics costs were to drop significantly. Our study confirms the potential of omics to predict additive and non-additive genetic effects, resulting in a potentially high increase in predictive abilities compared to standard genomic prediction models.

**Key Message:** Transcriptomics and proteomics information collected on a platform can predict additive and non-additive effects for platform traits and additive effects for field traits.

## 1 Introduction

Climate change affects crop production in several ways, such as significant changes in rainfall and weather patterns, increasing temperature, and drought stress (Lee et al. 2023). Such factors are threatening global food security, requiring immediate action to adapt crops to these challenging conditions better. The ever-decreasing cost of high-throughput phenotyping and genotyping technologies offers unprecedented opportunities to accelerate genetic gain by enhancing the efficiency of crop breeding programs. Especially in plant breeding, the interest in multi-environmental genomic predictions is constantly increasing thanks to pedo-climatic data availability (Westhues et al. 2022), which can help to model genotype by environment (GxE) interactions with greater predictive abilities (Crossa et al. 2022; Heslot et al. 2014; Jarquín et al. 2014; Rincent et al. 2019). Such environmental data, also known as environmental covariates (EC), can be included in different ways in multi-environment prediction models. For instance, the reaction norm approach by Jarquín et al. (2014) models markers and environmental covariates using covariance structures, where their interactions can be modeled through a Kronecker product between the genomic and the environmental covariance matrices. This model relies on a linear statistical relationship between genomic markers and the phenotype.

The advances in multi-omics technologies now make it possible to develop high-dimensional transcriptomic, proteomic, and metabolomic profiles that can efficiently capture the influence of environmental conditions on plants. Most studies that used metabolic data from field plants have proposed several approaches to integrate these in prediction models and have exhibited intermediate to high predictive abilities (de Abreu e Lima et al. 2017; Riedelsheimer et al. 2012; Ward et al. 2015), generally competitive with those obtained when using only genomic markers. Multi-omics data (Hu et al. 2021) are still expensive and cannot be made available for each trial of a breeding program. Breeders generally rely on phenotyping platforms to accurately simulate reproducible stressing conditions to study the multi-omics response of plants. Such data are quite helpful in identifying genes and QTLs related to stress responses (Blein-Nicolas et al. 2020; Jiang et al. 2024; Kilasi et al. 2018; Wen et al. 2019). However, their integration in the genomic prediction models of field traits is still quite challenging due to differences in environmental conditions in the field and platform.

Hu et al. (2020) illustrated that metabolomics and transcriptomics collected from the platform and the field are highly correlated between the same tissue types. This evidence suggests that platform omics data can be helpful for field predictions. It was the case in Fu et al. (2012) with platform transcriptomic data resulting in promising predictive abilities across genetically distant groups. Other studies have combined different platform omics, such as mRNA and sRNA transcriptomics and metabolomics data measured at an early stage, to improve the predictive abilities of field traits in maize (Schrag et al. 2018). For example, Westhues et al. (2017) collected transcriptomic data from plants in highly controlled conditions and reported improving their prediction models with transcriptomic data and genomic information while predicting maize field traits such as dry matter yield and dry matter content.

As omics abundances are influenced by the plant’s response to the growth conditions, we can expect them to be particularly efficient in predicting traits measured on the same plants or plants grown in similar conditions. Omics are intermediary phenotypes, so we hypothesize that they may efficiently capture non-additive genetic effects for predicting growth conditions like those in which the omics were measured. However, in contrast to DNA markers such as SNP, the heritability of omics measurements is lower than one. Moreover, Christensen et al. (2021) stressed that omics can capture genetic and non-genetic effects when the same individuals are characterized for omics and are predicted. The model proposed by Christensen et al. (2021), GOBLUP, can trim environmental noise and residuals from the omics expressions. Using this approach to remove the non-genetic part of the omics may be a way to increase the predictive ability of field traits with platform omics at the cost of removing the non-additive genetic part (i.e., the genetic part that is not captured by the additive kinship) of the omics.

Integrating such multi-faceted data can be done by fitting a multivariate linear mixed model (MvLMM). MvLMMs utilize correlations between traits to improve predictions. However, these correlations are not robust due to more predictors than the number of genotypes (p>>n) (Bernardo 2010). Further, the phenotypic correlations between traits are generally a poor representation of genetic correlations. Fitting such MvLMMs is quite computationally inefficient because of the number of iteratively estimated covariance components and their inverses, whose burden increases manifold with the increasing number of traits (Zhou and Stephens 2014). Runcie et al. (2021) proposed the Mega-Scale Linear Mixed Model (MegaLMM) as one possible solution that decreases the computational burden of MvLMM by re-parameterizing them as Bayesian sparse factor models. Like Bayesian Sparse Factor Analysis of Genetic Covariance Matrices (BSFG) (Runcie and Mukherjee 2013), MegaLMM also considers that a few latent factors can explain covariance among many traits. Following such an approach, MegaLMM has exhibited remarkable predictive abilities and runtime improvements over well-established Bayesian methods like Phenix (Dahl et al. 2016) and MTG2 (Lee and Van der Werf 2016) in wheat by using hyperspectral reflectance data and in *Arabidopsis thaliana* by using gene expression profiles (Runcie et al. 2021). MegaLMM is certainly a model of choice for predicting agronomic traits with multi-omics data.

In the present study, we worked on a panel of 244 maize Dent lines crossed to a Flint tester, characterized for ecophysiological traits, transcriptomics, and proteomics on a high-throughput phenotyping platform in two contrasted water treatments. In addition, grain yield was measured for this same panel of hybrids in 25 trials in different locations, years, and treatment combinations. Our primary objective was to assess the ability of multi-omics data obtained from platform experiments to predict both platform and field traits, in comparison to genomic selection. We worked both on platform and field traits, to evaluate the potential of platform omics to predict non-additive effects for traits measured in similar conditions (platform) or in completely different conditions (fields).

Additionally, we aim to evaluate their effectiveness in capturing GxE interactions in the fields. It involved a comparative analysis between models incorporating multi-omics data and conventional genomic prediction models relying solely on genomic information. Our particular focus lies in evaluating the potential advantages of employing the high-dimensional multivariate MegaLMM model in conjunction with omics data. Furthermore, we also assessed models combining both multi-omics and ECs.

## 2 Material and Methods

### 2.1 Plant material and experiments

#### 2.1.1 Maize panel and Platform experiments

The platform experiments were conducted on the PhenoArch platform (Cabrera‐Bosquet et al. 2016). The platform experiments comprised a hybrid panel of 254 maize dent lines crossed to a common flint parent (UH007) and grown under two contrasting water conditions, i.e., well-watered (WW) and water deficient (WD). The detailed experimental and growing conditions are available in Alvarez-Prado et al. (2018) and Blein-Nicolas et al. (2020). Experiments with two contrasting conditions were carried out in Spring 2012 and 2013 and Winter 2013. Plants grown in WW conditions faced a soil water potential of −0.05 MPa, while the ones grown under WD conditions faced a range from −0.3 to −0.6 MPa. Eight ecophysiological traits related to growth and transpiration were also measured during these experiments. The traits included early biomass (Be), late biomass (Biol), early leaf area (LAe), late leaf area (LAl), water use efficiency (WUE), water use (WU), stomatal conductance (gs), and transpiration rate (Trate). For each of these traits, we have up to three experiments within each treatment (Table S1).

#### 2.1.2 DROPS multi-environment trial

Phenology and production traits, including grain yield, were measured on 254 hybrids of the previously mentioned panel at 25 locations from Eastern to Western Europe and an external site in Graneros, Chile. For each experiment, genotypic means were computed for each hybrid using a mixed model with hybrids and replicates as fixed effects and spatial effects and spatially correlated errors as random effects (Millet et al. 2016). The model was fitted using ASReml-R (Butler et al. 2009; R Core Team 2013), and the best linear unbiased estimators (BLUEs) were used for the subsequent analyses. Like the platform experiments, DROPS trials were also conducted in two a priori contrasting water regimes, including WW and WD. ECs expected to affect maize growth at different phenological stages were measured in these trials, while some of them were computed using the APSIM crop growth model (Hammer et al. 2010; Millet et al. 2019). The covariates include longitude and latitude, soil water potential, maximum temperature, night temperature, radiation intercepted, and vapor pressure deficit during early flowering and grain-filling stages. A detailed overview of the environmental covariates is provided by Millet et al. (2019).

### 2.2 Multi-Omics Data

#### 2.2.1 Genomic Data

All lines were genotyped using the 50K Infinium HD Illumina array (Ganal et al. 2011), a 600K Axiom Affymetrix array (Unterseer et al. 2014), and more than 500K markers obtained through genotype by sequencing method (Negro et al. 2019). As a result, we had access to a dense panel of 978,134 markers spread across the maize genome. In addition, we applied a filter on minor allele frequency (MAF) and removed markers with an MAF lower than 0.02, resulting in 842,165 SNP markers to work with.

#### 2.2.2 Proteins

We had access to data on the abundances of 1,982 proteins from 254 hybrids grown under two contrasting watering conditions on the PhenoArch platform in the Spring 2012 experiment (Blein-Nicolas et al. 2020; Prado et al. 2018). For each hybrid, there were two replicates per condition, resulting in four leaf samples (one per plant) obtained at the pre-flowering stage. Proteins were extracted from the frozen leaves through standard trichloroacetic and acetone solutions-based methods and digested into peptides before mass spectrometry analysis. The proteins were quantified based either on extracted ion currents (XIC) or spectral count (SC). We used normalized and spatially corrected protein abundances as obtained from Blein-Nicolas et al. (2020). The broad sense heritabilities of the protein abundance adjusted means over replicates were computed independently for WD and WW treatments (Blein-Nicolas et al. 2020). For the analyses in this study, we filtered out proteins with a heritability lower than 0.4. We worked with 546 WD proteins and 475 WW proteins for trait predictions. Further details on protein extraction, quantification, spatial correction, and normalization can be found in Blein-Nicolas et al. (2020).

#### 2.2.3 ‘3’RNA seq Gene Expression Analysis

RNA was extracted from mature leaf tissues on the 1,524 plants from 254 different genotypes (254 hybrids x 2 treatments x 2 to 3 repetitions) of the spring 2013 platform experiment. The quantity of RNA was determined by spectrophotometry (Nanodrop (R), Ozyme), and the quality was evaluated by capillary electrophoresis (Agilent (R)) using the RNA Integrity Number (RIN). The concentration of RNA varied between 160 and 1000 ng/ml, and the RIN between 6.5 and 7. The NGS libraries were synthesized by using the QuantSeq-3’ mRNA-Seq Library Prep Kit FWD (Lexogen, Vienna). The quality and the quantity of each library were assessed using microfluidic electrophoresis on a LabChip GX (Perkin Elmer, Waltham, MA). 1,056 samples were selected among these 1,524 samples based on RNA quality (>200ng). For each sequencing batch, 1,056 samples were randomized between and within 11 plates of 96 wells according to four factors: genotype, treatment, genetic group origin, sampling date and hour, and line of sampling in a glasshouse using the function “designRandomize” R package (Brien, 2018). Accordingly, treatments, genetic structure, and sampling hour/date were well distributed between and within each plate. Then, 96 libraries of each sample in each plate were equimolarly pooled and sequenced on a Next Seq 550 (Illumina, San Diego, USA) using a High-output flow cell and a single read of 75 bases according to the manufacturer’s recommendations. The sequences were first trimmed with Cutadapt v1.8 on single read mode, removing adapter sequences and quality trimming (q 10). The minimum length to keep a read was 50. All samples were individually barcoded during the library process. 80% of the samples had more than 4 million reads, and 92% had more than 3 million reads. Alignment of the sequences was done with the software STAR, using reference genome v4 of B73 maize. The alignment was improved using an annotation file of the genome, allowing us to consider the splicing junctions. Only reads aligning to a single gene were considered. Moreover, only genes with at least one count per million reads in at least 10% of the samples were considered, which resulted in 20,475 and 20,642 transcripts for WD and WW conditions, respectively. In each condition, the transcripts were normalized with the approach Trimmed Mean of M-values (Robinson et al., 2010) and transformed with the log-cpm approach with package edgeR (McCarthy et al. 2012; Robinson et al. 2009).

For a given transcript, the last step was to fit a spatial model to compute adjusted means and estimate heritabilities. For this, we used the following model for each treatment (WW and WD):

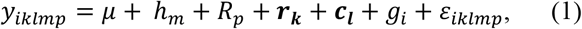

where *y*_*iklmp*_ is the observed value for transcript expression of hybrid *i* sown in row *k*, column *l*, repetition *p. h*_*m*_ is the fixed effect of the hour of sampling *m, R*_*p*_ is the fixed effect of repetition *p*, ***r***_***k***_ and ***c***_***l***_ are the random effects of row *k* and column *l*, respectively, *g*_*i*_ is the fixed genotypic effect of hybrid *i*, and *ε*_*iklmp*_ is the random error effect: 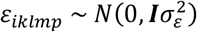. Here, ***I*** is the identity matrix and 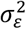 is the random error variance.

The SPaTS package (Rodriguez-Alvarez et al. 2018) within package statgenSTA (van Rossum et al. 2023) was used for this purpose, allowing for P-spline adjustment of the spatial effects. The number of segments used for adjusting the splines was 20 for rows and columns. The model was first fitted with a fixed genotype effect to compute marginal genetic means and then with a random genotype effect (identity matrix as variance/covariance matrix) to estimate broad sense heritabilities, as in Oakey et al. (2006). After the normalization procedure, and merging all the phenotypic and omics data, we obtained a list of 244 genotypes with transcript quantitative expression. For all further analyses, we filtered the transcripts based on their broad sense heritabilities to keep only those with heritabilities higher than 0.4. The filter resulted in 7,051 WD transcripts, 7,043 WW transcripts, 546 WD proteins, and 475 WW proteins.

### 2.3 Models

#### 2.3.1 Univariate Prediction Models

These univariate models were used for platform traits and field trait predictions.

##### GBLUP

As a reference, we applied a univariate GBLUP model for single trait or environment analysis, defined as follows:

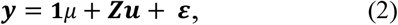

where ***y*** is the vector of adjusted means, **1** is the vector of 1s, *μ* is the overall mean. ***Z*** is the incidence matrix of random genetic effects, ***u*** is the vector of random genetic effects that are considered to follow a normal distribution, i.e., 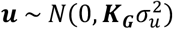. ***ε*** is the random error effect considered independent from ***u*** and following a normal distribution, i.e., 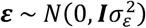. Here, ***K***_**G**_ is the genomic kinship matrix computed as ***K***_**G**_ = ***MM***^’^/2 ∑ *p*_s_ (1 − *p*_s_) with ***M*** as the matrix of centered marker genotypes and *p*_s_ as the frequency of reference allele *s* (VanRaden 2008). 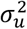 and 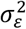 are the genetic and error variances, respectively. The GBLUP model was fitted using the R package rrBLUP (Endelman 2011).

##### OBLUP and G+OBLUP

Kernel-based methods generally offer an easy solution for multi-omics data integration. We used the following formula to calculate omics-based kernels:

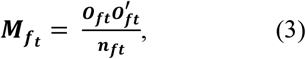

Where 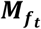 is the similarity matrix for omics type ***f*** in treatment ***t. O***_***ft***_ is the centered and scaled matrix of pretreated abundances of omics type ***f*** in treatment ***t***, and ***n***_***ft***_ is the number of features (proteins or transcripts) of the concerned type. All kernel matrices, including the kinship matrix, were scaled to have a sample variance of 1 (average of diagonal close to 1) to avoid biased parameter estimations due to different scales (Forni et al. 2011; Kang et al. 2010).

Firstly, we applied a univariate prediction model, hereafter referred to as OBLUP, with multiple random effects corresponding to different transcriptomic and proteomic effects as follows:

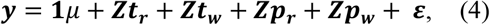

where **1**, *μ*, and ***ε*** are the same as defined earlier for equation 2. ***t***_***r***_ is the vector of random WD transcriptome-based genetic values; 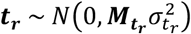. Similarly, ***t***_***w***_ is the vector of random WW transcriptome-based genetic values; 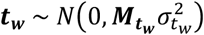. ***p***_***r***_ is the vector of random WD proteome-based genetic values; 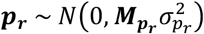. ***p***_**w**_ is the random WW proteome-based genetic values; 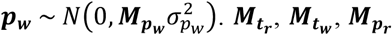 and 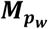 are the standard omics-based kernel matrices. All random effects are assumed to be independent.

Afterward, we extended the model to also include a standard random genetic effect (***u*)** as defined in equation 2. The extended model from hereafter will be referred to as G+OBLUP and can be represented as follows:

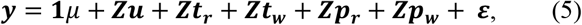

where **1**, *μ*, and ***ε*** are the same as defined in equation 2. ***t***_***r***_, ***t***_***w***_, ***p***_***r***_, and ***p***_***r***_ are the same as defined in equation 4. Like GBLUP, we applied both OBLUP and G+OBLUP on each platform and field trait one by one. We applied these models using R package Sommer (Covarrubias-Pazaran 2016).

##### AOBLUP and G+AOBLUP

We proposed AOBLUP and G+AOBLUP as the extensions of the two previous models, OBLUP and G+OBLUP. In the first step, we extracted the additive component (BLUPs) of omics features by applying an independent GBLUP model to each feature (transcript or protein):

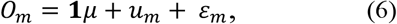

where *O*_*m*_ is the adjusted means of omics feature *m*, which can be either a single protein or transcript. *u*_*m*_ is the vector of random genetic effects of genome-wide markers on the omics feature and is considered to follow a normal distribution, i.e., 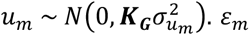 is the random error effect independent from *u*_*m*_ and follows a normal distribution as 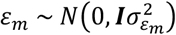. We also computed Pearson correlations between each omics as well as each additive component of omics as predicted by model (6) 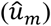, with phenotypic traits to evaluate which omics (standard or additive component) is the more useful for predicting a trait of interest.

In the second step, we estimated omics-based kernel matrices following the same formula as equation 3 but with the GBLUP of omics features (i.e., the additive part of omics features : 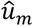) obtained in step one. In the third step, we fitted the AOBLUP and G+AOBLUP models separately by using additive omics-based covariance matrices, i.e.,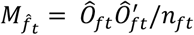, instead of standard omics-based kernel matrices. The AOBLUP model can be represented as follows:

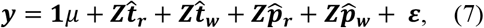

where **1**, *μ*, ***Z***, and *ε* are the same as defined earlier for equation 2. 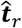 is the vector of random WD transcriptome-based additive genetic values; 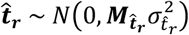. Similarly, 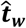 is the vector of random WW transcriptome-based additive genetic values; 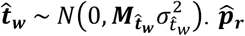 is the vector of random WD proteome-based additive genetic values; 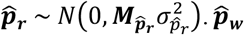 is the random WW proteome-based additive genetic values; 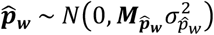. Here, 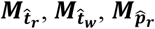, and 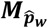 are additive omics-based kernel matrices.

While the G+AOBLUP can be represented as follows:

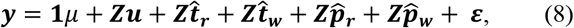

where **1**, *μ*, ***Z***, and ***ε*** are the same as defined in equation 2. 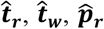, and 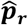 are the same as defined for the equation 7. Like GBLUP, we applied these models to all platform and field traits one by one. We applied these models using the R package Sommer (Covarrubias-Pazaran 2016).

## Multi-environment Prediction Models with Interactions for field traits prediction

Multi-environment prediction models enable information sharing between different environments, which can help in improving predictive abilities. These models were only used for field trait predictions.

### GxE-BLUP, GOxE-BLUP, and GAOxE-BLUP

The model, hereafter referred to as GxE-BLUP, integrates a GxE effect as follows:

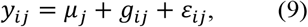

with *y*_ij_ being the phenotype of hybrid *i* in environment *j. μ*_*j*_ is the mean effect of environment *j. g*_*ij*_ is the random genetic effect of hybrid *i* in environment *j. ε*_*ij*_ is the residual of hybrid *i* in environment *j*. The random effect *g*_*ij*_ follows a normal distribution, i.e., *g*_*ij*_*∼MVR*(0, ∑), with ∑ = *K*_*G*_ ⊗ ∑_*E*_. Here, *K*_G_ is the genomic kinship matrix, ⊗ represents the Kronecker product, and ∑_*E*_ is the between-environment genetic covariance matrix with compound symmetry parametrization:

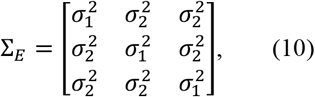

with 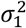 as the main genetic effect variance plus the GxE variance and 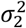 is the covariance between the environments. The compound symmetry model considers the same within environment variance and between environments covariance.

We introduce GOxE-BLUP and GAOxE-BLUP as extensions of GxE-BLUP to integrate the omics data. In these two models, we include both the main and interaction effects of omics. The first extension can be written as:

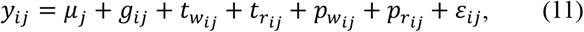

similar to *g*_*ij*_,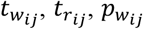 and 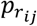 are the standard omics-based genetic effects of hybrid *i* in environment *j*. Like *g*_*ij*_, these effects also follow a multivariate normal distribution as; 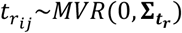, with 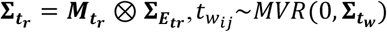with 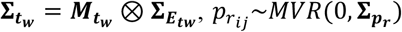, with 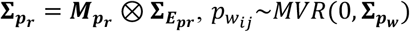 with 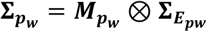. Here, 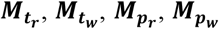 are the standard omics-based kinship matrices. 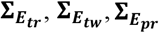, and 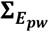 are the respective between environment genetic covariance matrices following compound symmetry parameterizations like **∑**_***E***_.

The second extension can be written as:

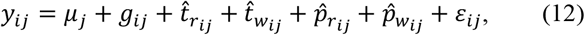

here, 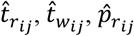, and 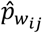 are the additive omics-based genetic effects of hybrid *i* in environment *j*. Like before in equation 11, these effects also follow a multivariate normal distribution as; 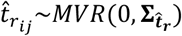, with 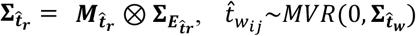, with 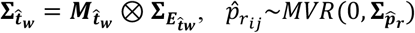, with 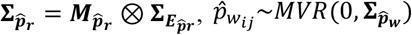, with 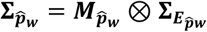. Here, 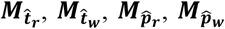 are the additive omics-based kernel matrices. 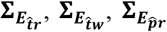, and 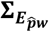 are the respective between environment genetic covariance matrices following compound symmetry parameterizations similar to **∑**_***E***_. We applied these models using R package Sommer (Covarrubias-Pazaran 2016).

### GxW-BLUP, GOxW-BLUP, and GAOxW-BLUP

ECs can also be used to describe environmental conditions that can impact and interact with the genotypes (Jarquín et al. 2014). We fitted the following model to include the effect of standardized ECs on traits (Jarquín et al. 2014):

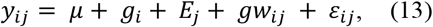

where *μ* is the overall mean, *g*_i_is the effect of genotype *i* and *E*_*j*_ is the effect of environment *j*. The term *gw*_*ij*_ is the genotype x ECs interaction term, which can be defined as 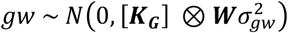, where, *Z*_*E*_ is the environmental incidence matrix, and 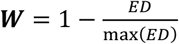. *ED* here represents the Euclidean distance between different environments calculated as 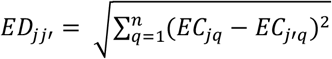 is the *q*th EC in the environment *j*.

Similar to GOxE-BLUP and GAOxE-BLUP, we extended the GxW-BLUP model into GOxW-BLUP (equation 14) and GAOxW-BLUP (equation 15) to account for standard and additive omics data, respectively.

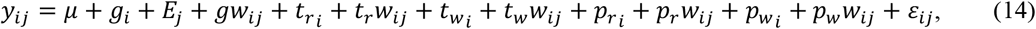

and

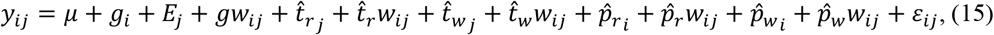

where *t*_r_*w*_*ij*_, *t*_w_*w*_*ij*_, *p*_r_*w*_*ij*_, and *p*_wr_*w*_*ij*_are the interactions between different standard omics-based genetic values and ECs, which can be defined as 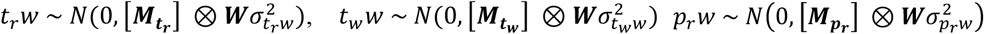,and 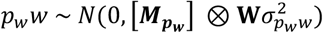, respectively. While,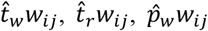, and 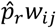 are the interactions between different additive omics-based genetic values and ECs, which can be defined as 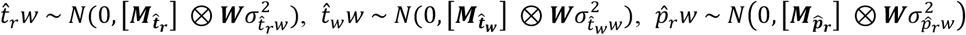 and 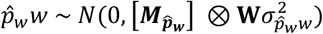, respectively. We fitted these models using the R package BGLR (Pérez and de los Campos 2014).

### MegaLMM

A classical MvLMM models the correlation between *t* traits through *t x t* genetic covariance (*G*_*m*_) and error (*R*) matrices. However, MegaLMM re-parametrizes MvLMMs as a factor model by introducing *K-* independent latent factors (Runcie et al. 2021). Unlike other models that use latent factors for dimension reduction and use factors to model a single random effect (mostly genetic) (Dahl et al. 2016; de Los Campos and Gianola 2007; Meyer 2007; Runcie and Mukherjee 2013), MegaLMM utilizes latent factors to model joint effects of all predictors, including fixed, random, and residuals. MegaLMM decomposes trait variation into two components, the first dependent on the *K* latent factors, which also model the correlations between the *t* traits and the second trait-specific variation uncorrelated between the traits. In other words, MegaLMM decomposes the variation of *t* traits into correlated and uncorrelated components. Further details about MegaLMM can be found in (Runcie et al. 2021). The final model implemented in this study can be written as follows:

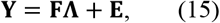

With

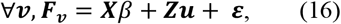

where **Y** is the *n x t* matrix for *n* genotypes and *t* traits. **F** is a *n x K* matrix of *n* genotypes and *K* latent factors (*K* is defined by the user). **Λ** is the *K x t* loading matrix whose elements, i.e. *λ*_*kj*_, determines the relation between the trait *k* and the corresponding factor *j*. **E** is a matrix of independent residuals for each trait. The *K* latent factors present in **F**, i.e., *F*_v_ for latent factor *v*, are further individually decomposed into fixed effect *β* and random effect ***u***, with ***X*** and ***Z***, respectively, as their incidence matrices. ***u*** is modeled as follows: 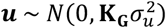. **ε** is the residual of each independent factor modeled as 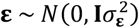.

The MegaLMM model uses Bayesian priors for regularization. We used Automatic Relevance Determination (ARD) (Mbuvha et al. 2020) for the factor loadings matrix. In high-dimensional multi-omics datasets with many features, ARD priors can help identify and emphasize important features, improving predictive ability and model interpretability. They introduce sparsity to the model by setting many predictor coefficients close to zero, aiding in feature selection and preventing overfitting. ARD priors are particularly useful when dealing with collinearity and limited sample sizes, as they enhance data efficiency and model stability. We fitted the MegaLMM models using the MegaLMM R-package (Runcie et al. 2021).

### Prediction scenarios

We compared two prediction scenarios that plant breeders classically face to evaluate the efficiency of the different models and omics data. The first one corresponds to classical situation in which breeders predict the performance of hybrids unobserved for the target trait (CV1). The second one predicts the performance of partially observed hybrids (CV2), as would be done in sparse testing. For both scenarios, all hybrids (training and test) had omics characterization (Fig. 1). Moreover, we used the same 5-fold and 5-repeat divisions of hybrids for all platform traits and grain yield trials for both CV1 and CV2. Comparing the predictive abilities of the different models for platform traits and field traits, allows us to evaluate the efficiency of omics data to predict non-additive effects when predicting traits measured in similar conditions (platform traits in a different experiment) or in completely different conditions (field) than those in which the omics were measured (the same platform experiment).

**Fig. 1.**
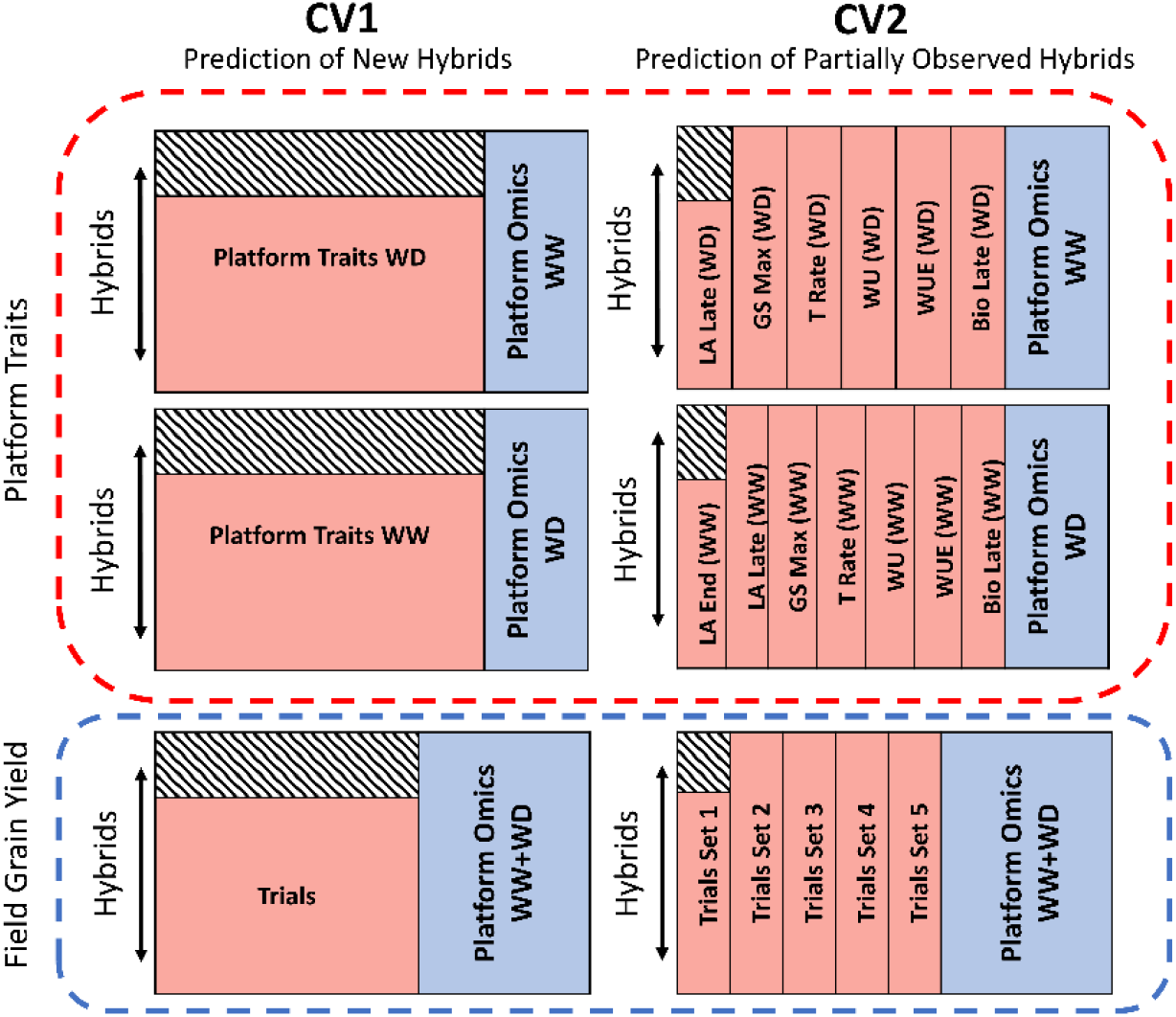
Description of the 5-fold and 5-repeat cross-validation schemes. CV1 is the scenario where we predict hybrids having no phenotypic observations but only platform omics data. CV2 is the scenario where we predict hybrids with partially observed phenotypes and complete platform omics data. The red (dashed) square represents the cross-validation settings of platform traits, while the blue (dashed) square represents the cross-validation settings of field grain yield. The pink color represents phenotypes available for training the models (platform traits or field grain yield), and the blue represents platform omics used as predictors. The hatched rectangle represents the hybrids/traits that are to be predicted.

### CV1 for Platform Traits

In CV1, we performed 5-fold with 5-repeats cross-validations for all the platform traits (Fig. 1). Meaning that for each iteration of the cross-validation, 20% of the hybrids were masked for all the platform traits and constituted the test set. This scheme was applied with MegaLMM, MegaGO, and MegaGAO. The omics information of both the training and test hybrids were used in models capitalizing on omics.

### CV2 for Platform Traits

In the CV2 scheme, we first categorized platform traits into trait sets. Each set contained different combinations of years and seasons for the same trait and the same water treatment. We formed six distinct trait sets for the WD platform traits, corresponding to traits LAe, gs, Trate, WU, WUE, and Biol, with three year-season combinations (three experiments) each. We created seven distinct trait sets for the WW platform traits, where the first two sets comprised one and two year-season combinations of LAe and LAl, respectively. While the remaining 5 comprised three year-season combinations of 5 platform traits: gs, Trate, WU, WUE, and Biol (Fig. 1). We performed 5-fold and 5-repeat cross-validation for each trait set and pooled all predictive abilities. Meaning that for each iteration of the cross-validation, 20% of the hybrids were masked for the trait set that had to be predicted. We applied this scheme to MegaLMM, MegaGO, and MegaGAO, with the last two also using omics information as predictors.

### CV1 for Field Traits (Grain Yield)

In CV1, we applied 5-fold cross-validation with 5 repeats, in which we masked the phenotypes of the test hybrids across all the field trials (Fig. 1). We applied this validation scheme with the multienvironment and multivariate models defined above, i.e., GxE-BLUP, GxW-BLUP, MegaLMM, GOxE-BLUP, GOxW-BLUP, MegaGO, GAOxE-BLUP, GAOxW-BLUP, and MegaGAO. We also included WW and WD omics information for both training and test hybrids for the models capitalizing on omics.

### CV2 for Field Traits (Grain Yield)

In CV2, we first randomly assigned the 25 yield trials in five sets of five trials each, referred to as Trial Sets 1 to We performed 5-fold and 5-repeat cross-validation for each trial set (Fig. 1). It means that for each iteration of the cross-validation, 20% of the hybrids were masked for the trial set that had to be predicted. We applied this validation scheme with the same multivariate models mentioned in the previous section. We also included omics information for both training and test hybrids for the models capitalizing on omics.

As univariate models (GBLUP, OBLUP, G+OBLUP, AOBLUP, and G+AOBLUP) consider a single platform trait or a grain yield trial at a time, they resulted in the same predictive abilities in CV1 and in CV2. In addition, models that included a GxE component or ECs were only applied to grain yield field trials.

An important point to note is that, unlike grain yield, we independently predicted the two platform trait treatments (WW and WD). We predicted WW platform traits with omics originating from WD treatment and predicted WD platform traits with omics originating from WW treatment (Fig. 1). We implemented this precaution to ensure that the predictive capabilities based on omics solely depended on genetic effects rather than any potential non-genetic effects that might arise from obtaining omics data and traits from the same plants.

As this study relies on model comparison, we used gain to measure model performance compared to the reference. We defined gain as the predictive ability of each model minus the ability of univariate GBLUP, which only uses information from within the predicted trial or platform trait, as was the case for all univariate models. Predictive ability was defined as the Pearson correlation between the predictions and the adjusted means of the test hybrids.

## Results

### Univariate and multivariate models for platform trait prediction without transcript and protein information

#### CV1 Predictions

We predicted platform traits with different univariate and multivariate models under CV1 (Table 1 & Fig. 2). Firstly, in the case of WD traits, the reference univariate GBLUP model resulted in a mean predictive ability of 0.26 over all the traits. The multivariate MegaLMM model resulted in a predictive ability of 0.25 with negligible difference compared to the reference GBLUP model. The results of WW platform traits are similar to the WD platform traits. For instance, the reference GBLUP model resulted in a mean predictive ability of 0.24, while the multivariate MegaLMM model resulted in 0.25.

**Table 1.**
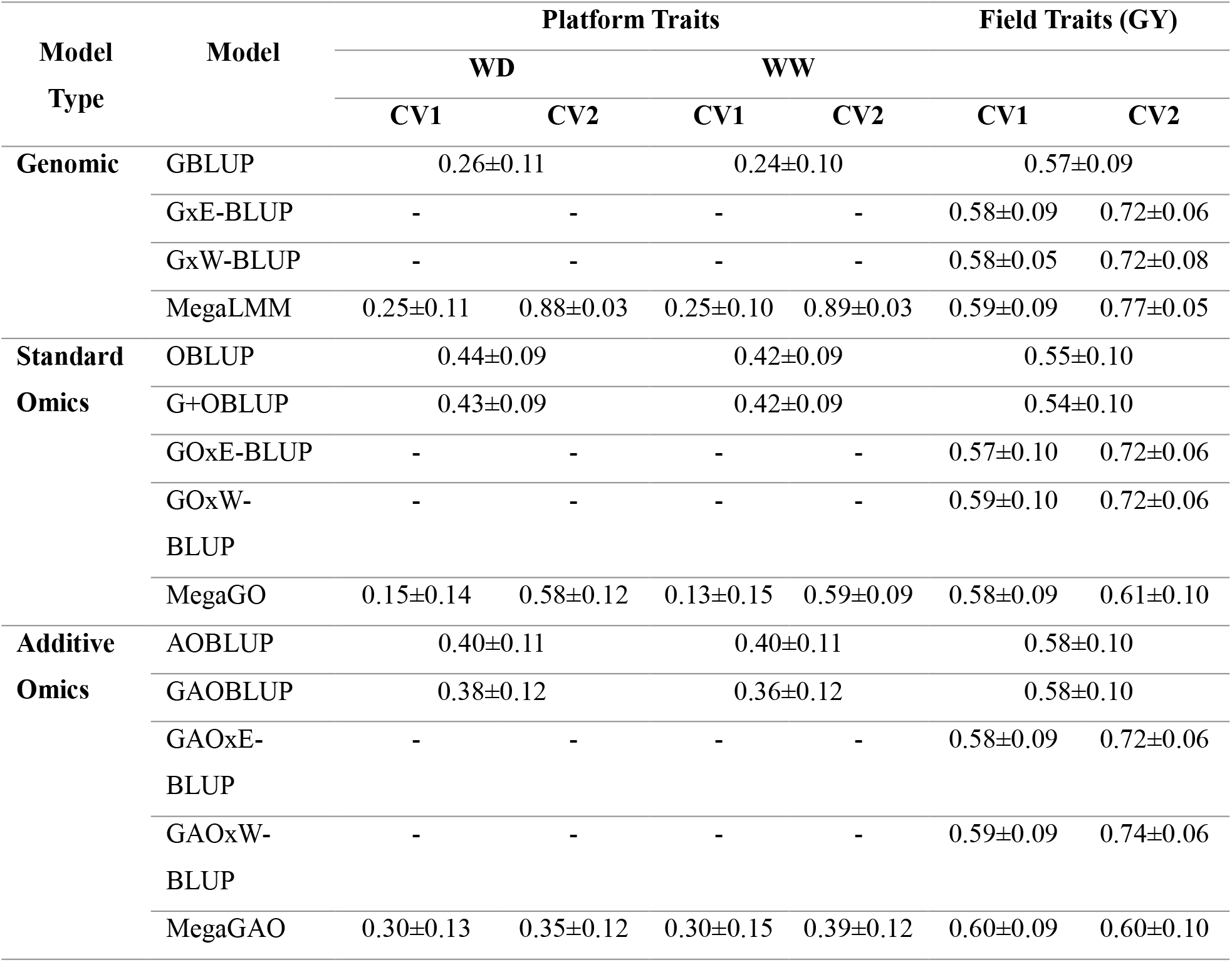
Predictive abilities of different genomic and omics models for predicting WD & WW platform traits and field traits (grain yield) in CV1 and CV2 cross-validation schemes. Model type represents whether the model has genomic, standard omics, or additive omics information. Univariate models presented were tested only trait by trait for platform and environment by environment for grain yield under 5-fold-5-repeat CV, resulting in the same predictive abilities for CV1 and CV2. Standard deviations are computed individually for each platform trait and grain yield, followed by a mean over platform traits or field trials.

**Fig. 2.**
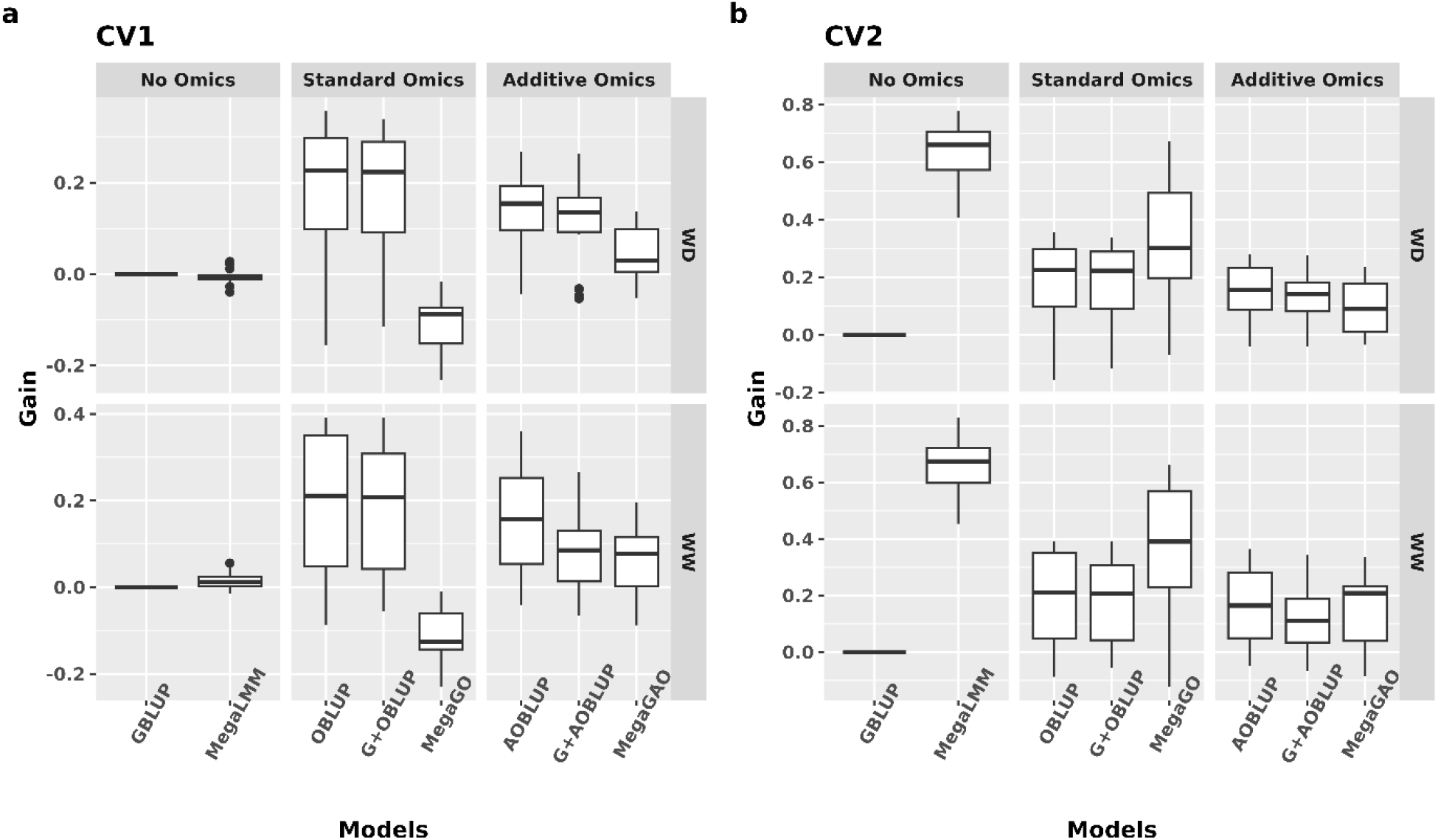
Boxplots of predictive gains of different models compared to GBLUP in different cross-validation schemes for platform traits. A) CV1 Scheme for predicting new hybrids, B) CV2 Scheme for predicting partially observed hybrids. Here, WD represents WD platform traits predicted with WW omics, and WW represents WW traits predicted with WD omics. “No Omics” represents the scenarios without transcripts and proteins, “Standard Omics” represents the case where standard transcripts and proteins are used, and “Additive Omics” represents the case where the additive component of transcripts and proteins was used.

#### CV2 Predictions

In the case of CV2 (Table 1 & Fig. 2) with WD traits, the MegaLMM model resulted in a predictive ability of 0.88, 0.62 units higher than the reference GBLUP. Similar results were also observed with WW platform traits in CV2 with MegaLMM, resulting in a mean predictive ability of 0.89.

### Univariate and multivariate models for platform trait prediction with transcript and protein information

#### CV1 Predictions

We used WW omics as predictors for WD platform traits (Table 1 & Fig. 2). The OBLUP model applied to WD traits resulted in a mean predictive ability of 0.44, which is 0.18 units higher than the benchmark GBLUP model. Adding genomic information in addition to transcripts and proteins, i.e., in the G+OBLUP model, we observed a similar mean predictive ability of 0.43. We observed a slight decrease in predictive abilities when we shifted to additive omics. The model solely with transcripts and proteins as predictors, i.e., AOBLUP, resulted in a mean predictive ability of 0.40. At the same time, the model that also used genomic information (G+AOBLUP) resulted in a mean predictive ability of 0.38.

The predictive abilities of models predicting WW platform traits were similar to those of WD platform traits. We used WD omics as predictors for WW platform traits (Table 1 & Fig. 2). The OBLUP model resulted in a mean predictive ability of 0.42. Adding genomic information (G+OBLUP) did not change the predictive ability. As before, we observed a decrease in mean predictive abilities with additive omics. For example, the AOBLUP model resulted in a predictive ability of 0.40, while G+AOBLUP resulted in an even lower predictive ability of 0.36.

Now, we will compare different multivariate prediction models with omics information for predicting platform traits (Table 1 & Fig. 2). In the case of WD traits, MegaGO, the model with standard omics, resulted in a mean predictive ability of 0.15, lower than any other prediction model for the same traits. Shifting to additive omics with MegaGAO resulted in a mean predictive ability of 0.30, higher than MegaGO but quite lower than all other univariate models for the WD platform traits.

The performances of multivariate MegaGO and MegaGAO for predicting WW platform traits (Table 1 & Fig. 2) were similar to their respective cases with WD platform traits. For example, both resulted in mean predictive abilities of 0.13 and 0.30, respectively, again falling behind univariate models for the same traits.

#### CV2 Predictions

Under CV2, we tested two models capitalizing on omics information, MegaGO and MegaGAO (Table 1 & Fig. 2), with the former utilizing standard omics while the latter utilizing additive omics. The MegaGO model applied to WD platform traits resulted in a mean predictive ability of 0.58, indicating a gain of 0.32 units over the reference GBLUP model but a loss of 0.30 units compared to MegaLMM, the model without omics. Shifting to additive omics made the case worse. MegaGAO resulted in a mean predictive ability of 0.35, comparable to the reference GBLUP and lower than MegaLMM and MegaGO. We also applied MegaGO and MegaGAO models to WW platform predictions (Table 1 & Fig. 2). Both models performed similarly as in the predictions of WD platform traits, resulting in mean predictive abilities of 0.59 and 0.39, respectively.

### Univariate and multivariate models for field trait predictions without transcript and protein information

#### CV1 Predictions

At first, we applied both univariate and multivariate models to predict grain yield without adding omics information (Table 1 & Fig. 3). The reference GBLUP model resulted in a mean predictive ability of 0.57 over all the folds and repeats of all 25 field trials. The two interactions-based models, i.e., GxE-BLUP and GxW-BLUP, resulted in similar mean predictive abilities of 0.59 and 0.58, respectively. Similarly, the multivariate MegaLMM model also resulted in a mean predictive ability of 0.59. Even though the average predictive abilities of MegaLMM appear to be closer to the reference model, there are several trials where important gains were observed, for instance, 0.06 points in the case of “Karlsruhe.2012.OPT” in contrast to the minimum gain equal to 0 in “Campagnola.2012.WD” Fig. S1).

**Fig. 3.**
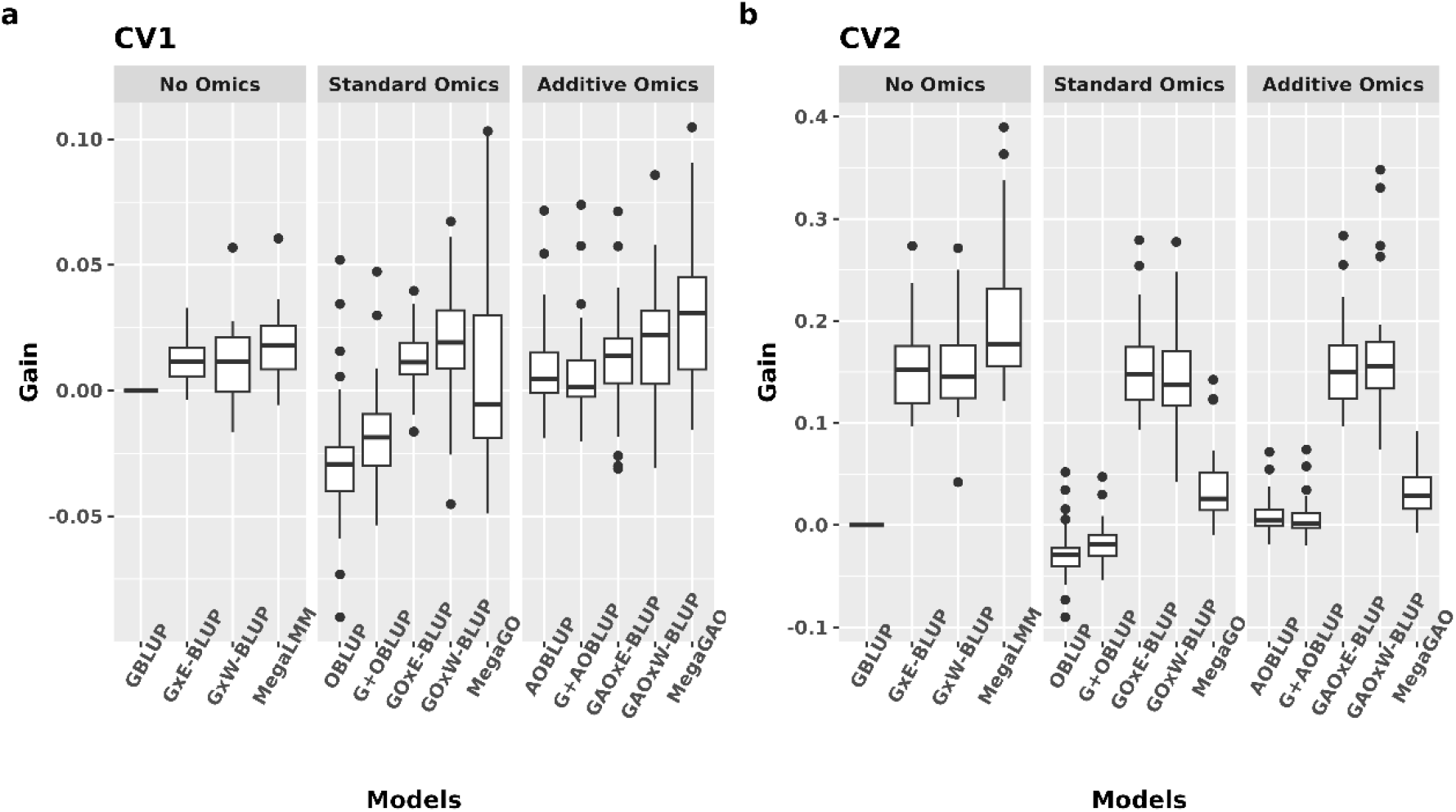
Boxplots of predictive gains of different models compared to GBLUP in different cross-validation schemes for field grain yield. a) CV1 Scheme for predicting new hybrids, b) CV2 Scheme for predicting partially observed hybrids, “No Omics” represents the scenarios without transcripts and proteins, “Standard Omics” represents the case where standard transcripts and proteins are used, and “Additive Omics” represents the case where the additive components of transcripts and proteins were used.

#### CV2 Predictions

We tested GxE-BLUP, GxW-BLUP, and MegaLMM in a less challenging CV2 situation for predicting grain yield (Table 1 & Fig. 3). In this cross-validation scenario, GxE-BLUP and GxW-BLUP models resulted in a similar predictive ability of 0.72, which is 0.15 units higher than GBLUP. MegaLMM performed better than the reference GBLUP and the two interaction-based models, with a mean predictive ability of 0.77. We also observed considerable gains with MegaLMM for many environments. For instance, “Craiova.2012.WD” improved by 0.39 units in comparison to GBLUP (Fig. S2).

### Univariate and multivariate models for field trait predictions with transcript and protein information

#### CV1 Predictions

Firstly, we tested the predictive ability of the omics data to predict field traits one by one using univariate models (Table 1 & Fig. 3). We observed that adding standard omics information in the univariate models decreased the predictive abilities compared to GBLUP (Table 1 & Fig. 3). For example, OBLUP and G+OBLUP resulted in mean predictive abilities of 0.55 and 0.54, respectively, lower than the GBLUP benchmark model (0.57). Shifting to additive omics with AOBLUP and G+AOBLUP models, we observed the same predictive ability of 0.58 with both models. Even though their predictive ability is somewhat higher than OBLUP and G+OBLUP, it is similar to that of the reference GBLUP.

The two interaction-based models with standard omics, GOxE-BLUP and GOxW-BLUP, resulted in predictive abilities of 0.57 and 0.59, respectively, similar to their predecessors, GxE-BLUP and GxW-BLUP. The two interaction-based models with additive omics, GAOxE-BLUP and GAOxW-BLUP, resulted in comparable predictive abilities of 0.58 and 0.59, respectively. Even though the performance is similar, GAOxW-BLUP resulted in important predictive gains in some trials, as high as 0.09 (Craiova.2012.WD), with a maximum loss of −0.03 (Campagnola.2013.OPT), as observed in Fig. 4.

**Fig. 4.**
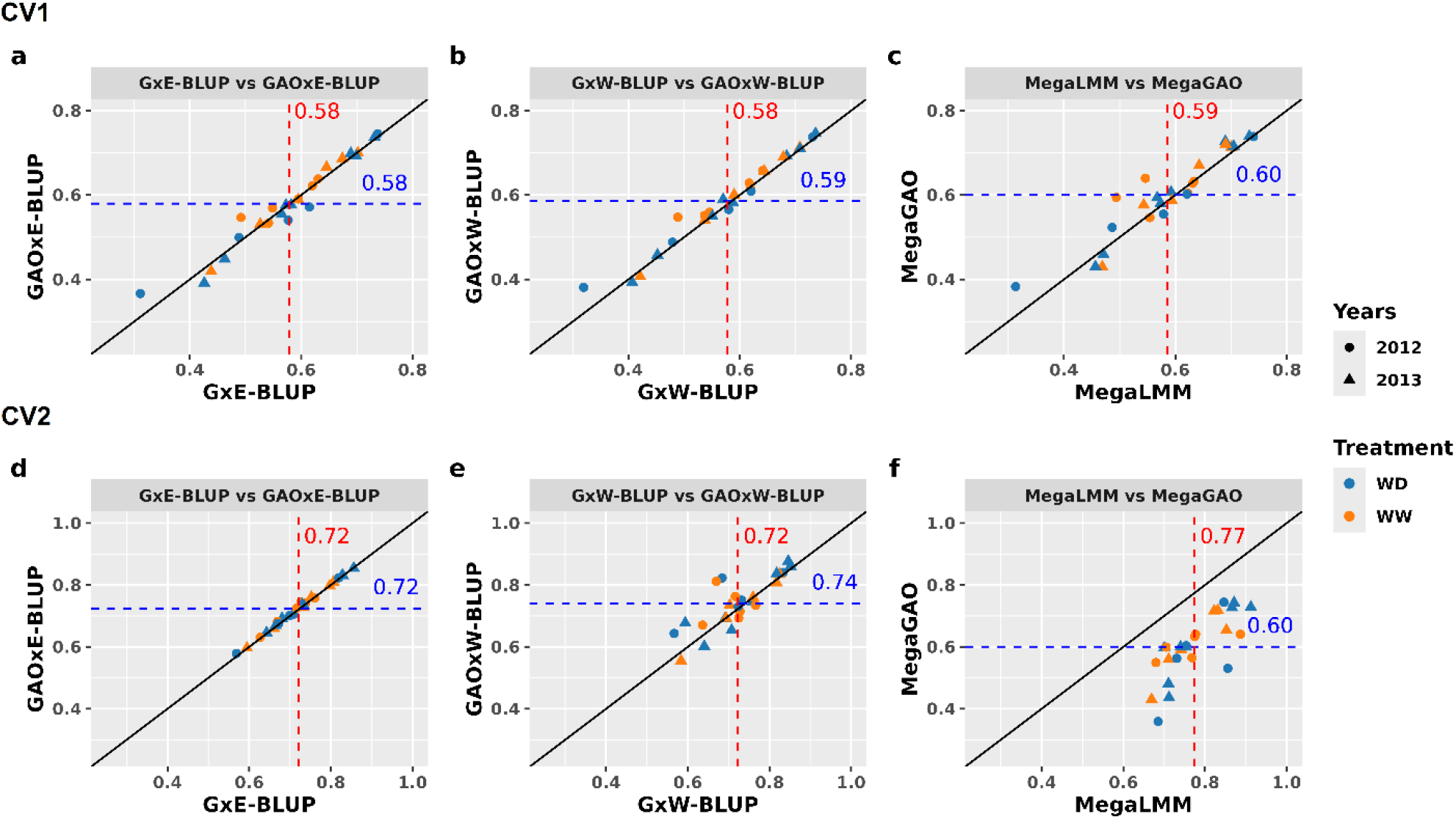
Comparison of predictions of different additive omics-based models with their versions without omics in CV1 (top row) and CV2 (bottom row) for grain yield. The X-axis represents the model without omics, and the Y-axis represents the model involving additive omics. The red dashed line represents the mean predictive ability of the model without omics (x-axis), while the blue dashed line represents the mean predictive ability of the model involving additive omics (y-axis). The black line represents the diagonal. Colors and the shapes of points refer to the treatment and years of field trials.

The predictive ability of multivariate MegaGO (0.58) was also quite similar to the reference GBLUP and MegaLMM models. However, using additive omics information, MegaGAO showed some improvement over GBLUP and MegaLMM with a mean predictive ability of 0.60. Even though we could not observe overall important improvements by using additive omics data in MegaGAO, some environments, such as “Craoiva.2012.OPT” and “Campagnola.2012.OPT”, indicated gains as high as 0.10 compared to MegaLMM, while the maximum loss was 0.04 in “Campagnola.2013.OPT” (Fig. 4). Overall, in the challenging situation of CV1, all the models, including the additive omics information, exhibited gains relative to their predecessors with standard omics information, although these gains were small.

#### CV2 Predictions

The interaction-based models involving standard omics, GOxE-BLUP and GOxW-BLUP, resulted in the same predictive abilities (0.72) as GxE-BLUP and GxW-BLUP, suggesting that omics information was not beneficial in the CV2 scenario. The additive omics-based models, GAOxE-BLUP and GAOxW-BLUP, resulted in mean predictive abilities of 0.72 and 0.74, respectively. Even though the gain with GAOxW-BLUP compared to GxW-BLUP is small, there are several trials where we observed gains as high as 0.09 (Craiova.2012.WD) and a maximum loss of −0.03 (Campagnola.2013.OPT) (Fig. 4). Multivariate MegaGO with standard omics information resulted in a mean predictive ability of 0.61, indicating a sharp decrease compared to 0.77 of MegaLMM (Table 1 & Fig. 3). It suggests that adding standard omics in the model was worsening the situation. This situation is also true with the multivariate MegaGAO model containing additive omics information.

## Discussion

The interest in utilizing omics data in trait predictions is increasing due to their ability to capture interactions and regulation processes in response to the environment. However, due to the high costs of multi-omics characterization, it is currently impossible for a breeder to collect such data for all field plots each year. It is possible to measure omics in controlled conditions on phenotyping platforms. Even though omics data originating from the platform help detect stress QTLs (Blein-Nicolas et al. 2020), their capabilities to predict platform and field traits need further evaluation. Therefore, we evaluated different univariate and multivariate models with or without omics data. Our main objectives were to assess the efficiency of those omics and of a high dimensional multivariate model, MegaLMM (Runcie et al. 2021), to predict traits measured in similar conditions (platform), or to predict productivity traits measured in completely different conditions (the fields), than those in which the omics were measured (a different platform experiment). For this, we have considered standard univariate models such as GBLUP, OBLUP, AOBLUP, G+OBLUP, and G+AOBLUP, predicting each platform trait or each grain yield trial one by one. We also fitted GxE-BLUP and GxW-BLUP models, that were developed to capture GxE in multi-environment settings. We extended these models to be able to include standard omics, GOxE-BLUP, and GOxW-BLUP, or additive omics, GAOxE-BLUP and GAOxW-BLUP. We have compared the efficiency of the multi-trait models in two different prediction schemes, CV1 and CV2. CV1 focuses on predicting new hybrids that are not phenotyped for platform or field traits but have multi-omics characterization. CV2 predicts hybrids partially observed for some platform traits or field trials. CV2 is generally easier to predict than CV1 due to the presence of performances of test hybrids in unmasked platform traits or field trials.

### Platform Omics are very efficient in predicting additive and non-additive genetic effects for platform traits

We first tested and compared models without using the omics information to predict platform traits. In the case of CV1, MegaLMM performs similarly to GBLUP for both WW and WD platform traits. It is consistent with studies showing that multi-trait models are not much better than univariate models if none of the traits is phenotyped for the test set. In the case of CV2, MegaLMM performed extremely well (0.77) compared to GBLUP (0.57, Table 1). In this case, the MegaLMM model benefits from the presence of different correlated phenotypes of test hybrids for unmasked traits. This scenario, similar to trait-assisted prediction, has been shown to result in highly accurate predictions when predicted and observed traits are correlated (Calus and Veerkamp 2011; Fernandes et al. 2018; Henderson and Quaas 1976; Jia and Jannink 2012; Pszczola et al. 2013; Robert et al. 2020).

We also tested different omics and additive omics-based prediction models on platform traits. OBLUP and G+OBLUP, the two univariate prediction models with standard omics information were much better than the reference GBLUP model (Table 1 & Fig. 2), with gain in predictive ability as high as 0.36 (Spring_2013.WD.WU) and 0.34 (Spring_2012.WD.WU), respectively. It means that omics are very efficient to make predictions in environmental conditions similar to those in which they were measured. This is of particular interest for prediction approaches based on physiological traits measured in controlled conditions (Bouidghaghen et al. 2023) Note that we ensured that the improved predictive abilities of models OBLUP and G+OBLUP were only due to genetics (and not to sharing some environmental factors) by predicting traits measured on different plants than those used for omics measurements. More precisely, we used omics measured in the WD treatment to predict plants grown under the WW treatment and the reverse. This means that the only non-genetic effect that could be involved is the quality of the seed lots, as the same seed lots were used for both treatments.

One key result is that upon fitting the models with additive omics, we observed an important decrease in predictive abilities with AOBLUP and G+AOBLUP compared to OBLUP and G+OBLUP. This means that the non-additive genetic part of the omics was very useful in predicting platform traits. Another important result to note is that omics measured in similar conditions as the predicted trait (but different water treatments) can capture some non-additive genetic variation like epistasis and responses to growth conditions. We can indeed hypothesize that responses to local conditions are driven by regulatory processes, with epistasis as a consequence. These findings are also in line with the suggestions of Li et al. (2019) regarding the importance of omics expressions in capturing non-additive genetic variation. The potential ability of OBLUP to capture non-additive genetics works when conditions are similar because the epistatic variation is likely correlated more than between platform and field.

### GxE-BLUP and GxW-BLUP models are useful for predicting grain yield

The benchmark GBLUP model (**Erreur ! Source du renvoi introuvable. &** Fig. 3) resulted in a mean predictive ability of 0.57 for grain yield. In both CV1 and CV2 validation schemes, multi-environment models GxE-BLUP and GxW-BLUP models performed better than GBLUP. As expected, the gains with models in CV1 are not as prominent as in CV2. These results are comparable to Burgueño et al. (2012), where authors observed a significant improvement in CV2 but not in CV1. Even though our GxE-BLUP model performs extremely well, it has one disadvantage due to its underlying assumption of the same variance and covariance among different trials. Furthermore, we expected the model with envirotyping data, i.e., GxW-BLUP, to perform better than GxE-BLUP, but this was not the case. The potential advantage of using GxW-BLUP is to give more weight to trials with similar environmental conditions. Literature indicates that including environmental covariates is not always advantageous. For instance, Monteverde et al. (2019); Nguyen et al. (2023) reported that adding environmental covariates resulted in worse predictions than GBLUP in rice while predicting new environments, which could be attributed to higher inter-environmental differences. In the studies predicting new genotypes, Jarquín et al. (2014); Malosetti et al. (2016) and Rincent et al. (2019) observed little to no improvement with a GxW model.

### MegaLMM is an efficient model to predict field traits in multi-environment scenarios

We also implemented a two-level hierarchical model, MegaLMM (Runcie et al. 2021). MegaLMM was the best model for both CV1 and CV2 prediction schemes. The gain was marginal for CV1, but MegaLMM seemed to be very efficient at dissecting genetic correlations between trials and can benefit from those, particularly in the CV2 scenario, with a gain of 0.21 units in comparison to GBLUP and 0.05 compared to GxE-BLUP and GxW-BLUP. In previous studies, MegaLMM was accordingly shown to result in a high predictive gain compared to GBLUP in sparse testing scenarios (Runcie et al. 2021). However, it was not tested in the more challenging situation of CV1 before. These results confirm that multi-trait models are very helpful when some information on the predicted set is available, for example, in the case of trait-assisted predictions or sparse trial networks.

### The inclusion of platform-based omics can only slightly improve the predictive ability of field traits

We also fitted different prediction models with standard omics or with the additive part of the omics. We hypothesized that extracting the additive genetic component of omics might be more beneficial for field predictions, because it is a way of removing part of the error (Christensen et al. 2021), and because non-additive genetic effects are probably different on the platform than in the field, and so not useful for field trait predictions. It is clearly visible in Fig. 5 that the additive omics expressions have a higher Pearson correlation with grain yield than the standard omics, which was not the case for platform traits (Fig. S3). In CV1 (Fig. 3), none of the models sufficiently benefited from adding the standard omics. However, with the addition of additive omics instead of standard omics, most of the models exhibited small to intermediate gains. For instance, the AO-BLUP model appeared to be 0.03 units better than OBLUP but similar to GBLUP, meaning that the additive part of omics alone can predict similarly to genomic markers. It is likely that additive omics-based kernel matrices get closer to the genomic kinship matrix, which is evident from the higher correlations between additive omics and genomic kinship matrices (0.68, 0.61, 0.62, and 0.60 for 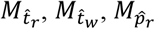, and 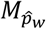, respectively) compared to standard omics and genomic kinship matrices (0.59, 0.56, 0.58, and 0.57 for 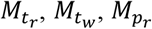, and 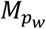, respectively). The AO-BLUP model does not include genomic markers in the regression step on field traits but includes them in the first step, where we remove non-additive genetic and residual noise from the omics. Even though gains with omics information were low on average, predictions of some environments showed important improvements, whereas just a few environments exhibited small losses (Fig. 4).

**Fig. 5.**
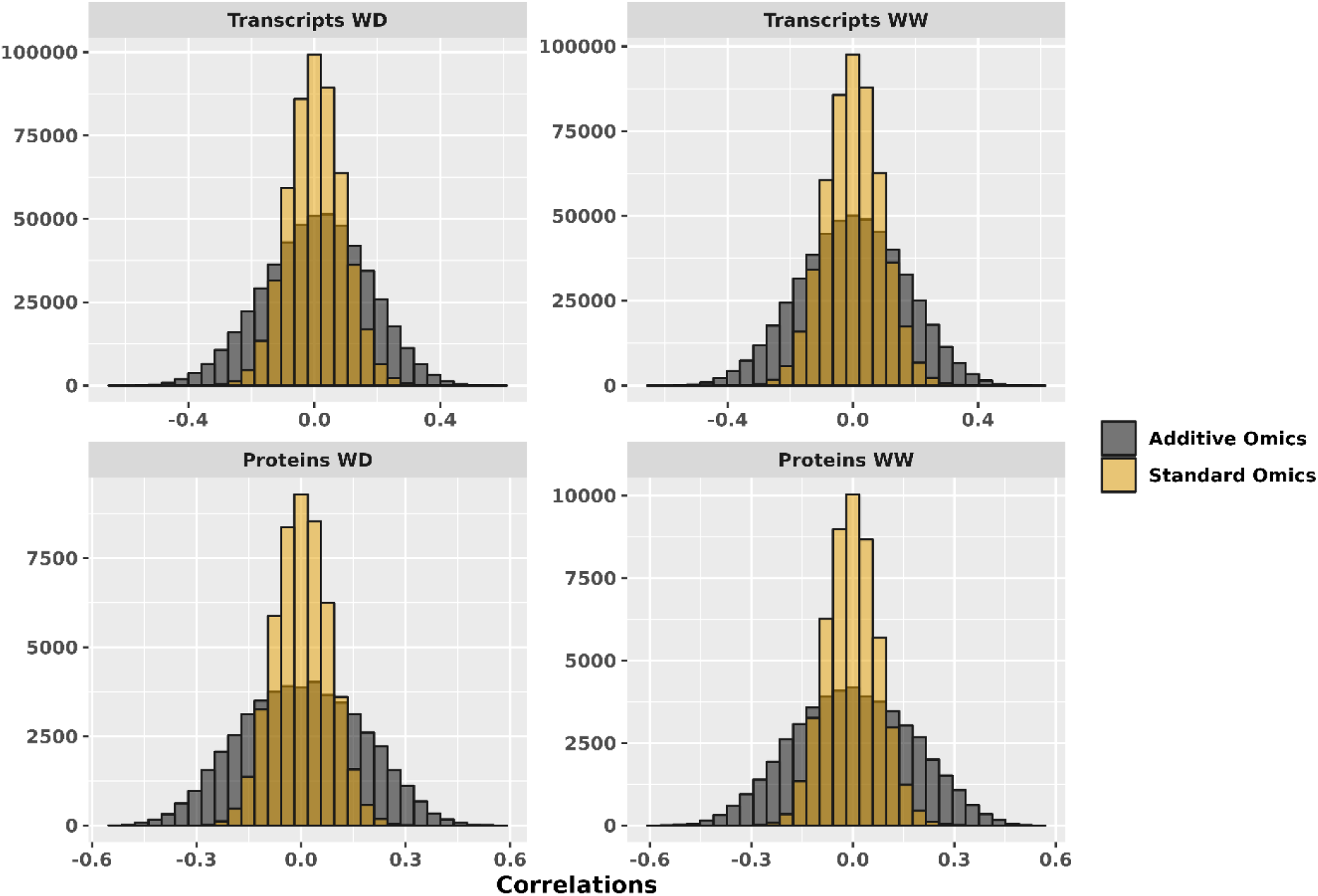
The distribution of the correlations between omics and grain yield from all 25 trials. The grey color refers to the additive component of omics, and the yellow color refers to the standard omics. WD represents omics measured in water-deficit and WW represents omics measured in well-watered conditions. The top row corresponds to transcripts and the bottom row corresponds to proteins.

As the omics capture a response to the environmental conditions, we also hypothesized that transcriptomics and proteomics could be useful to model GxE interactions, which are important in field trials as they can account for a significant part of phenotypic variance, sometimes even more than the genetic variance (Rogers et al. 2021). In our dataset, the amount of variance explained by GxE is similar to the variance explained by G alone (Millet et al. 2016). We proposed two multi-environment models with omics information, and according to our knowledge, our approach to model interactions between omics and environmental covariates in a multi-environment setting is the first implementation in the literature. In the CV1 scheme, the GAOxW-BLUP model gains slightly compared to the same model without omics (Table 1). The gain compared to GxW-BLUP is higher in CV2, where GxW-BLUP predictive ability stands at 0.72 and GAOxW-BLUP at 0.74, with some environments gaining as high as 0.14 units (Fig. 4). If we also compare these results to GxE-BLUP and GAOxE-BLUP, the improvement with GAOxW-BLUP might likely have occurred due to the interactions between additive omics and ECs.

For CV1, the best model was the one based on additive part of omics and MegaLMM, i.e., MegaGAO, which was expected since trimming off residual variance and noise from the omics increased correlations with the grain yield (Fig. 5). Upon removing the non-additive part from the omics data, we observed a strong increase in the number of features with an absolute correlation above 0.4. Overall, additive omics are useful in CV1 because they offer some phenotypic information about masked hybrids, which is better than having no information at all. The performance of MegaGAO in CV2 was lower than expected. MegaGAO loses in CV2 despite unmasked field trials probably because their information gets diluted within the high dimensions of omics information, and the quality of information provided by the omics is lower than that of unmasked trials. This case of MegaGAO in CV2 was also true for MegaGO.

The scientific community’s interest in implementing biological information in genomic prediction models to boost genetic gain is consistently increasing (MacLeod et al. 2016). Our work indicates that omics data can be considerably useful for making predictions in conditions similar to those in which they were measured. When predicting field traits with platform omics, the gain was only marginal and only the additive part of the omics was useful. We conclude from this, that measuring the omics in field conditions is necessary to capture non-additive effects and boost predictive ability. It was for instance proposed that near-infrared data measured in the fields could be useful for predicting GxE interaction (Robert et al. 2022a; Robert et al. 2022b).This may also be possible with molecular omics in the near future thanks to the important decrease of cost that are ongoing.

## Conclusion

In this study, we integrated omics information collected from the platform to predict field and platform traits using different univariate and multi-environmental models and a mega-scale multivariate model, i.e., MegaLMM. We also introduced a novel multi-environmental model to integrate omics and ECs simultaneously (GOxW-BLUP). Platform omics data could lead to dramatic increase in predictive ability compared to GBLUP for traits measured in similar conditions, but the gain was marginal for predicting traits in completely different conditions (platform versus fields). The main reason was that platform omics were able to predict non-additive effects only for plants grown in similar conditions. In this case, the non-additive component of omics becomes important due to its ability to capture non-additive genetic variations like epistasis related to the response to the environmental conditions. For field traits, we observed that trimming omics data off their non-additive genetic and residual components can improve field predictions, especially while predicting new hybrids, because such components cannot explain the field phenotypic variation. We also observed that MegaLMM could extract information from additive omics in predicting field traits, particularly for predicting new hybrids, which were only evaluated for omics information. Finally, in the absence of omics, we confirm the very high predictive abilities of multi-trait model MegaLMM in the sparse testing scenario (CV2).

## Author Contribution Statement

AB performed the statistical modelling and analyses under the supervision of RR, DR, AC, TM-H and LM. AB wrote the manuscript. RR edited the manuscript. DR supported AB for the MegaLMM analyses. LCM, SN, FT and HD designed the ‘3’RNAseq experiment. HD pretreated and mapped the ‘3’RNAseq data, BH-B and ML treated and analyzed the ‘3’RNAseq data under the supervision of SN, RR and FC. CW and FC coordinated the field and platform experiments. All authors approved the final version of the manuscript for submission.

## Data and script Availability

Field grain yield and environmental covariates can be found in: https://doi.org/10.1104/pp.16.00621. Platform phenotypic data are available online using the PHIS information system (Neveu et al. 2019) at http://www.phis.inra.fr/openphis/web/index.php?r=site%2Flogin-as-guest. From this webpage, data are accessible by clicking on Experimental organization > Project > Systems genetics for maize drought tolerance (Amaizing project). For proteomics data, The raw MS output files were submitted to PROTICdb (http://moulon.inra.fr/protic/amaizing) (Ferry-Dumazet et al. 2005; Langella et al. 2007, 2013), the MassIVE database (https://massive.ucsd.edu/ProteoSAFe/static/massive.jsp) under accession number MSV000085594, and the ProteomeXchange database (http://www.proteomexchange.org) under accession number PXD019804. Detailed information on all peptides and proteins identified in the LC-MS/MS runs, as well as peptide intensities and protein abundances obtained for each sample, are also freely available on PROTICdb at the same URL. The genotyping data can be found at: https://doi.org/10.15454/AEC4BN. Transcriptomics data are available to anyone upon reasonable request. R scripts to run the analysis are available at: https://github.com/rrincent/MegaGO.

## Declaration

The authors declare no conflict of interest or personal relationships that could appear to influence the work reported in this paper.

## Acknowledgments

We thank all the collaborators and fundings of the PIA Amaizing (ANR-10-BTBR-01) and the DROPS (FP7-244374) projects for the production of these very rich datasets. We are grateful to the INRAE MIGALE bioinformatics facility (MIGALE, INRAE, 2020. Migale bioinformatics Facility, doi:10.15454/1.5572390655343293E12) for providing help and/or computing and/or storage resources. We thank Melisande Blein-Nicolas, Michel Zivy and the platform PAPPSO for the introduction to the world of proteomics.

