## Supplementary figures for "High-dimensional multi-omics measured in controlled conditions are useful for maize platform and field trait predictions"

### Supplementary Tables

**Table S1** List of platform traits and their short names and counts in WW and WD Treatments. The WW and WD treatments were applied one next to the other in the same three experiments.

| Platform Trait | Short Name | Number of experiments (WD treatment) | Number of experiments (WW Treatment) |
| --- | --- | --- | --- |
| Late Leaf Area | LAl | 3 | 2 |
| End Leaf Area | LAend | 0 | 1 |
| Maximum Stomatal Conductance | gs | 3 | 3 |
| Transpiration Rate | Trate | 3 | 3 |
| Water Use | WU | 3 | 3 |
| Water Use Efficiency | WUE | 3 | 3 |
| Late Biomass | Biol | 3 | 3 |

### Supplementary Figures

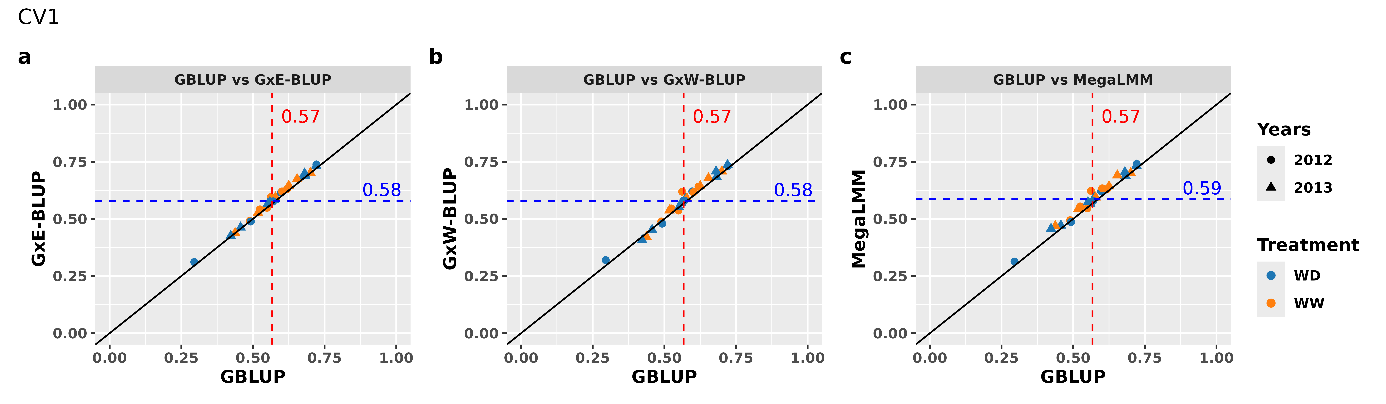

**Fig. S1** Comparison of predictions of different models with GBLUP in CV1 for grain yield. The X-axis represents the reference GBLUP model, and the Y-axis represents the model under consideration. The red dashed line represents the mean predictive ability of GBLUP, while the blue dashed lines represent the mean predictive ability of the model under consideration. The black line represents the diagonal. Colors and the shapes of points refer to the treatment and years of field trials. a) GxE-BLUP, b) GxW-BLUP, and c) MegaLMM.

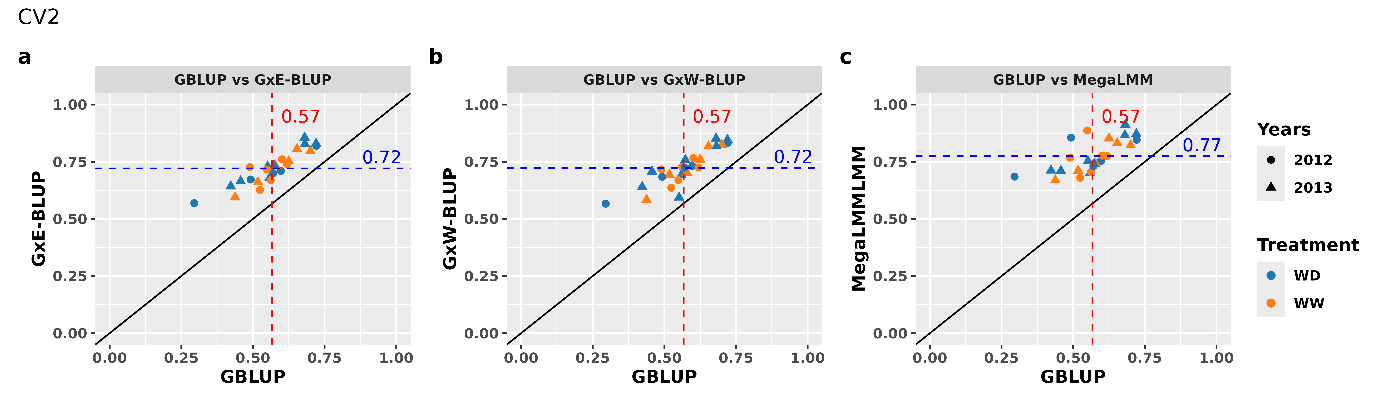

**Fig. S2** Comparison of predictions of different models with GBLUP in CV2 for grain yield. The X-axis represents the reference GBLUP model, and the Y-axis represents the model under consideration. The red dashed line represents the mean predictive ability of GBLUP, while the blue dashed lines represent the mean predictive ability of the model under consideration. The black line represents the diagonal. Colors and the shapes of points refer to the treatment and years of field trials. a) GxE-BLUP, b) GxW-BLUP, and c) MegaLMM.

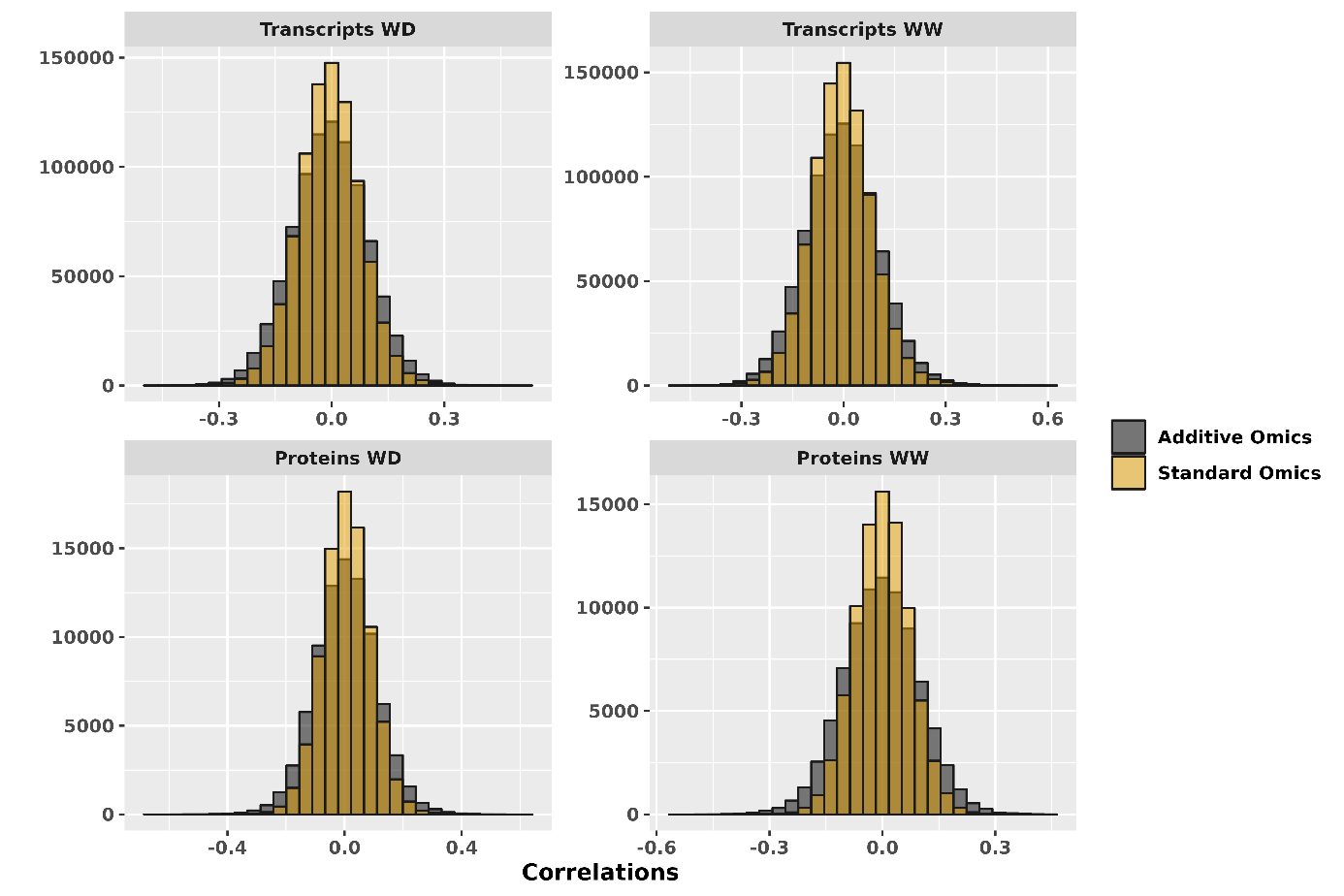

**Fig. S3** The distribution of the correlations between omics and platform traits. The pink color refers to the omics, and the cyan color refers to the additive components of omics. A) Transcripts from Well-Watered treatment, B) Transcripts from Water-Deficit treatment, C) Proteins from Well-Watered treatment, and D) Proteins from Water-Deficit treatment.
